# A tale of two fish: Comparative transcriptomics of resistant and susceptible steelhead following exposure to *Ceratonova shasta* highlights differences in parasite recognition

**DOI:** 10.1101/2020.06.04.133991

**Authors:** Damien Barrett, Jerri L. Bartholomew

## Abstract

Diseases caused by myxozoan parasites represent a significant threat to the health of salmonids in both the wild and aquaculture setting, and there are no effective therapeutants for their control. The myxozoan *Ceratonova shasta* is an intestinal parasite of salmonids that causes severe enteronecrosis and mortality. Most fish populations appear genetically fixed as resistant or susceptible to the parasite, offering an attractive model system for studying the immune response to myxozoans. We hypothesized that early recognition of the parasite is a critical factor driving resistance and that susceptible fish would have a delayed immune response. RNA-seq was used to identify genes that were differentially expressed in the gills and intestine during the early stages of *C. shasta* infection in both resistant and susceptible steelhead *(Oncorhynchus mykiss).* This revealed a downregulation of genes involved in the IFN-γ signaling pathway in the gills of both phenotypes. Despite this, resistant fish quickly contained the infection and several immune genes, including two innate immune receptors were upregulated. Susceptible fish, on the other hand, failed to control parasite proliferation and had no discernible immune response to the parasite, including a near-complete lack of differential gene expression in the intestine. Further sequencing of intestinal samples from susceptible fish during the middle and late stages of infection showed a vigorous yet ineffective immune response driven by IFN-γ, and massive differential expression of genes involved in cell adhesion and the extracellular matrix, which coincided with the breakdown of the intestinal structure. Our results suggest that the parasite may be suppressing the host’s immune system during the initial invasion, and that susceptible fish are unable to recognize the parasite invading the intestine or mount an effective immune response. These findings improve our understanding of myxozoan-host interactions while providing a set of putative resistance markers for future studies.

## Introduction

*Ceratonova shasta* (syn. *Ceratomyxa shasta)* is a myxozoan parasite of salmonid fish that is endemic to most river systems in the Pacific Northwest of the United States [1,2]. It is recognized as an economically important pathogen of both wild and hatchery-reared salmonids [3–6] and has been linked to population-level declines [7,8]. *C. shasta* has a broad host range and is able to infect most, if not all, native salmonid species [2]. The initial site of infection is the gills, where the parasite spore attaches to the epithelium prior to invading the blood vessels and beginning replication. Travelling via the bloodstream, it reaches the intestine 4 to 5 days after the initial infection, where it continues to replicate and undergoes sporogenesis [9]. Severe infections result in enteronecrosis (ceratomyxosis) and death of the host. Fish stocks in the Pacific Northwest are highly divergent in their innate resistance to *C. shasta* induced mortality: those originating from *C. shasta* endemic watersheds exhibit a high degree of resistance [8,10], whereas allopatric fish are highly susceptible [8,11]. Numerous studies have demonstrated that resistance to *C. shasta* is a genetically controlled trait that shows little variation within a given population [12–17].

While the innate resistance of the host is a primary factor in the outcome of infection, disease severity falls on a spectrum that is heavily influenced by the exposure dynamics, which include exposure concentration and duration, water temperature, and parasite virulence [8]. At the very low end of this spectrum, susceptible fish appear unable to mount an effective immune response to *C. shasta* and suffer mortality rates at or near 100% at doses as low as one spore per fish [11,18]. When resistant fish are exposed under similar conditions, few if any parasites reach the intestine and no clinical signs of disease are observed [19–21]. However, if the exposure dose is high, typically greater than 10,000 spores, resistant fish may succumb to the infection and the disease progresses as it does in susceptible fish [9,22]. When resistant fish experience more intermediate exposure conditions, *C. shasta* is observed reaching the intestine but the fish are able to control and eventually clear the infection [23]. Bartholomew et al. found that resistant steelhead *(Oncorhynchus mykiss)* and cutthroat trout *(O. clarkii)* chronically exposed to *C. shasta* at low temperatures (< 10° C) had infections characterized by large numbers of parasites on the intestinal mucosal surface and multiple foci of inflammation in that tissue [5]. However, sporogenesis was not observed, mortality rates were low, and observations of fibrosis in histological sections suggested that fish were recovering from the infection. Containment of the parasite in well-defined granulomas has also been observed in sub-lethal exposures of resistant steelhead trout and Chinook salmon (*O. tshawytscha)* [20,22,24].

Understanding the host response to infection is complicated by the fact that *C. shasta* is a species complex, comprised of four distinct genotypes that have different salmonid host associations: genotype 0 with *O. mykiss;* genotype I with Chinook salmon; and genotype II, which is considered a generalist that is able to infect multiple fish species but contains a mix of two genetically distinct subtypes named after their associated hosts: IIR for rainbow trout (freshwater strain of *O. mykiss)* and IIC for coho salmon (*O. kisutch)* [2,25–27]. Along with different host specificities, these genotypes have different effects on their hosts. Genotype 0 typically causes chronic infections with no apparent morbidity or mortality. In contrast, genotypes I and II may be highly pathogenic in their respective hosts, causing the disease signs that are classically associated with *C. shasta* infections.

Knowledge of the infecting genotypes, and establishment of parasite’s lifecycle in a laboratory setting [28], has permitted investigations of the immune response to *C. shasta* be conducted in a controlled setting with known genotypes. One of the first, by Bjork et al., compared the host response of susceptible and resistant Chinook salmon to *C. shasta* genotype I infection [25]. No difference in parasite burden at the gills was detected. However, in the intestine, resistant fish had both a lower infection intensity and a greater inflammatory response than susceptible fish and were able to eventually clear the infection. Both phenotypes had elevated expression of the pro-inflammatory cytokine IFN-γ in the intestine, but only susceptible fish had elevated levels of the anti-inflammatory cytokine IL-10. A similar trend was found in a subsequent study of susceptible rainbow trout infected with genotype IIR, with significant upregulation of IFN-γ, IL-10, and IL-6 [29]. It has also been demonstrated that fish exposed to *C. shasta* are able to produce parasite-specific IgM and IgT [30,31]. Both IgM and IgT were found to be upregulated in high mortality genotype IIR infections [29], but whether this antibody response offers any protection against *C. shasta* pathogenesis remains to be determined.

Currently, no prophylactic or therapeutic treatments exist for *C. shasta* induced enteronecrosis and efforts to manage the disease revolve around selective stocking of resistant fish. However, even resistant fish may succumb to infection [8] and assessing the resistance level of a fish stock requires a series of lethal parasite challenges with large groups of fish. Insight into the molecular and genetic basis of resistance will help facilitate the development of vaccines and therapeutics for this pathogen as well as provide a non-lethal biomarker for assessing a stock’s resistance. More broadly, the immune response to myxozoan pathogens remains largely uncharacterized, having been explored in a limited number of species. As a result, there is a near complete lack of therapeutics or other disease control measures, an issue that is becoming more evident as aquaculture continues to increase worldwide [32,33]. *C. shasta* genotype II presents a unique model for studying the immune response to myxozoans as it is highly virulent and fish hosts are either highly resistant, or completely susceptible to the parasite, rather than falling on a continuum. Additionally, the resistance phenotype of many fish stocks is already known, which avoids the issue of *ad hoc* determination of phenotype or the need to create resistance and susceptible lines of fish for research. *C. shasta* is also one of the few myxozoans whose complete life cycle is both known and maintained in a laboratory setting.

The fact that *O. mykiss* is the primary fish host is also advantageous, as rainbow trout is one of the most widely studied and cultivated fish species and an extensive knowledge base exists for it, including a fully sequenced genome. Taken together, we believe that the *C. shasta-O. mykiss* system offers a tractable model for studying the immune response to myxozoans and what genes drive resistance.

With this in mind, we chose to use resistant and susceptible steelhead as model for understanding how and when the host responds to infection at the transcriptomic level. We hypothesized that early recognition of the parasite by the host was a critical factor in resistance and that susceptible fish would fail to recognize the initial infection, responding only after the parasite began to proliferate within the intestine. Conversely, we hypothesized that resistant fish would quickly recognize and respond to the infection, preventing parasite establishment in the intestine and proliferation once there. To test this, we held both phenotypes in the same tank and exposed them in parallel to *C. shasta* to ensure equivalent exposure conditions. Infected tissue was collected from both phenotypes at 1, 7, 14, and 21 days post exposure (dpe) to assess parasite proliferation using qPCR (all timepoints) and the local host immune response during the early stages of infection (1 and 7 dpe) using RNA-Seq.

## Material and methods

### Fish

Resistant steelhead from the Round Butte Hatchery and susceptible steelhead from the Alsea Hatchery, both located in Oregon, USA, were used in this study. From each hatchery, 6 adults were collected (3 male, 3 female) and bred to create pure-parental offspring. The offspring were raised at the Oregon State University (OSU) John L. Fryer Aquatic Animal Health Laboratory in Corvallis, Oregon, USA. The fish were fed daily with a commercial diet (Bio-Oregon, Longview, Washington, USA), and reared in tanks supplied with 13.5° C specific-pathogen free (SPF) well water. Two weeks prior to the parasite challenge, the fish were fin-clipped for identification and transferred to 100-liter tanks and acclimated to 18°C. This temperature was chosen as it reflective of the river water temperatures that out-migrating salmon experience when they are exposed to *C. shasta*, and aligns with previous studies [34].

### Parasite challenge

*C. shasta* genotype IIR actinospores were collected from two colonies of *Manayunkia occidentalis,* the freshwater annelid host [35], which were maintained in indoor mesocosms receiving flow-through UV-treated river water. Influent water to each colony was shut off 24 hours prior to the challenge to allow actinospores to accumulate in the mesocosm water. To ensure that both the resistant and susceptible fish were exposed to the same concentration of actinospores, 50 fish (susceptible average 42.2 ± 3.2 g; resistant average 39.4 ± 2.9 g) from each stock (differentiated on the presence of a fin clip) were placed together in identical control and treatment tanks containing 375-liters of water maintained at 18°C. Three liters of mesocosm water, which contained an estimated 4,500 actinospores based on monitoring of parasite production by qPCR [36], was added to the treatment tank. At the same time, three liters of water from an uninfected annelid mesocosm was added to the control tank. Fish were held on static water with aeration for 24 hours, at which time each treatment group (resistant exposed, resistant control, susceptible exposed, susceptible control) was sorted and placed into triplicate 25-liter tanks (12 total) that were randomly assigned and supplied with 18°C water. Water samples were collected from the exposure tanks immediately after the mesocosm water was added and after the fish were removed to quantify the number of *C. shasta* spores present at the beginning and end of the challenge. The water samples were immediately filtered and prepared for qPCR following a previously described method [36].

### Sample collection

Fish were sampled at 1, 7, 14, and 21 days post exposure (dpe), with 1 dpe corresponding to 24 hours after initiation of parasite exposure. Fish were sampled at the same time of day to minimize possible changes in gene expression due to circadian rhythms [37]. At each timepoint, 3 fish from each tank were euthanized with an overdose of MS-222 (tricaine methanosulfonate, Argent Laboratories, Redmond, WA, USA) for a total of 12 fish per treatment group, and 48 per timepoint. From 2 of the 3 fish, gills (1 dpe) or intestine (7, 14, 21 dpe) were collected whole and immediately placed in RNAlater and stored at 4° C for 24 hours, prior to being placed at −80° C for long term storage. From the remaining fish, gills and intestine were collected and placed in Dietrich’s fixative for histology. All methods involving live fish were approved by Oregon State University’s IACUC (protocol # 4660). A summary diagram of the experimental setup in shown in Fig 1.

**Fig 1.**
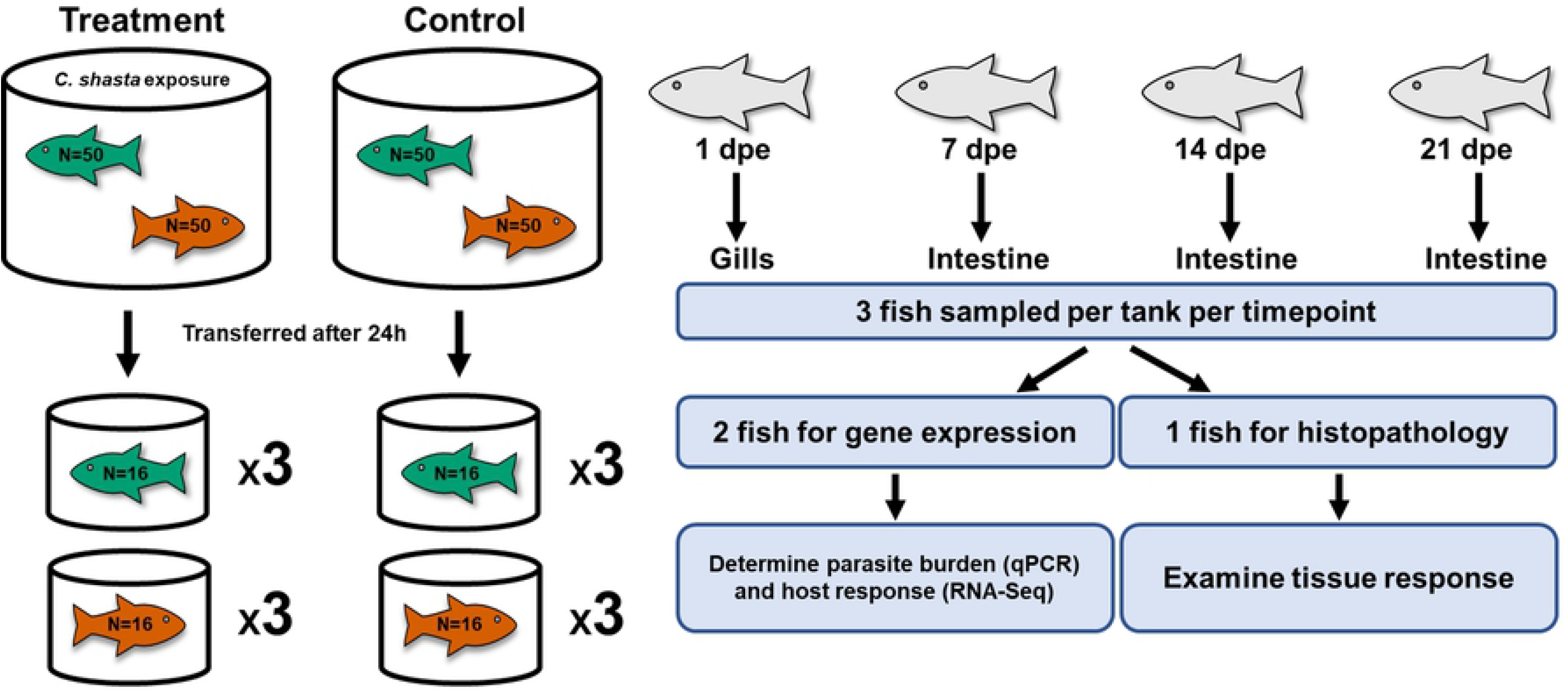
Experimental diagram of the exposure conditions and subsequent sampling of steelhead. Susceptible steelhead (green) and resistant steelhead (orange) were exposed to *Ceratonova shasta* for 24 hours and then each phenotype was separated and placed into triplicate tanks. Resistant fish had been previously fin-clipped as a means of identification. dpe = days post exposure.

### Sample processing

Due to variation in the size of the gills and intestine between fish, each tissue was homogenized in liquid nitrogen using a porcelain mortar & pestle and subsampled. RNA was extracted from 25 mg of homogenized tissue using the RNeasy Mini Kit (Qiagen, catalog number 74104) following the manufacture’s protocol. DNA was extracted from 25 mg of homogenized tissue from each sample using the DNeasy Blood & Tissue Kit (Qiagen, catalog number 69506) and eluted in 30 μl of Buffer AE, applied to the spin column twice, to achieve a higher concentration. The purity and concentration of the extracted RNA and DNA was assessed using a NanoDrop ND-1000 UV-Vis Spectrophotometer.

To assess the parasite load in each of the tissues, a previously developed *C. shasta* qPCR assay [36] was used to quantify the amount of parasite DNA present. 100 ng of DNA extracted from each sample was assayed in triplicate wells through 40 cycles using an Applied Biosystems StepOnePlus Real-Time PCR System. A sample was considered positive for *C. shasta* if all wells fluoresced and the sample was rerun if the Cq standard deviation between wells was greater than 1. On each qPCR plate, a positive control, a negative control (molecular grade water), and a standard curve of dilutions equivalent to 1, 10, 100, and 1000 actinospores was included.

Histological sections were prepared by the OSU Veterinary Diagnostic Laboratory, Corvallis, OR, USA and stained with H&E.

### Sequencing

To understand the transcriptomic response of both resistant and susceptible fish during the early stages of *C. shasta* infection, mRNA from the gills at 1 dpe and from the intestine at 7 dpe was chosen for sequencing. To control for possible confounding variables, such as tank effects, six samples from each treatment group were chosen at random and were evenly split across the three tanks housing each group. 48 samples (24 per timepoint) were submitted to the Center for Genome Research and Biocomputing at OSU for library preparation and sequencing. The integrity of the RNA was confirmed by running each sample on an Agilent Bioanalyzer 2100 (Agilent Technologies, USA). 1 ug of RNA was used for library preparation using the Illumina TruSeq™ Stranded mRNA LT Sample PrepKit according to the manufacturer’s instructions (Cat. No. RS-122-2101, Illumina Inc. San Diego, CA, USA). Library quality was checked with a 4200 TapeStation System (Agilent Technologies, USA) and quantified via qPCR. All libraries were sequenced on 4 lanes of an Illumina HiSeq 3000 as 100-bp single-end runs. The libraries were randomly distributed across the 4 lanes, 12 per lane.

Examination of the sequencing data from 7 dpe led us to sequence intestinal mRNA from susceptible fish at 14 and 21 dpe to follow the response in a progressive infection. Since we anticipated large differences in gene expression at these timepoints due to the intense histological changes observed, we chose to sequence six samples from each timepoint (3 exposed, 3 control) and do so at a higher depth of coverage to account for a greater proportion of the sequenced reads coming from parasite mRNA. 12 samples (6 per timepoint) were submitted for library preparation and sequencing as described above and were sequenced on two 100-bp single-end lanes. Resistant fish were not sequenced at these timepoints due to the low infection prevalence and intensity, the minimal transcriptomic response at 7 dpe, and because no tissue response was observed by histology.

### Data analysis

Adapter sequences were trimmed from the raw reads using BBDuk (January 25, 2018 release), which is part of the BBTools package [38], and all reads less than 30-bp after trimming were discarded. Library quality was assessed before and after trimming using FastQC (v0.11.8) [39]. Reads were then mapped to the latest rainbow trout reference genome (GenBank: MSJN00000000.1) using HiSat2 (v 2.1.0) [40]. Due to the high number of homeologs present in the *O. mykiss* genome [41], the aligned reads were filtered and sorted using SAMtools (v 1.9) [42] to exclude all reads that mapped to more than one location in the genome. The number of reads that mapped to each gene was calculated using HTSeq-count (v 0.11.1) [43] and the raw counts imported in R 3.4.1 [44] and loaded into the package DESeq2 (v 1.18.1) [45]. To identify potential outliers, heatmaps and PCA plots were constructed from the raw counts that were regularized log-transformed using the DESeq2 function rlogTransformation().

Differentially expressed genes (DEGs) were identified using the negative binomial Wald test in DESeq2 and were considered significant at a Benjamini–Hochberg False Discovery Rate (FDR) adjusted p-value < 0.05 and an absolute log_2_(fold change) > 1. Annotation of the DEGs and gene ontology (GO) enrichment was conducted with Blast2GO (v 5.2.5) [46] with a blast e-value cutoff of 1e^-5^. To obtain high quality and informative annotations, genes were preferentially annotated with the SWISS-PROT database [47] followed by the NCBI nonredundant database and a taxonomy filter of ‘Actinopterygii’ and ‘Vertebrata’ was applied. All genes detected within a tissue were used as the background for GO enrichment. Enriched GO terms along with their FDR-adjusted p-values, were imported into Cytoscope (v 3.7.2) [48] for visualization with the ClueGo (v 2.5.6) [49] plugin, which clusters genes and GO terms into functionally related networks. *O. mykiss* was chosen as the organism for Ontologies/Pathways and the GO Term Fusion option was used to merge GO terms based on similar associated genes. Volcano plots were constructed with the R package EnhancedVolcano (v 1.0.1) [50].

### RNA-seq validation by quantitative reverse transcription PCR (RT-qPCR)

The expression of four immune genes *(IFN-y, TNF-α, IL-10, IL-1β)* found to be differentially expressed by RNA-seq were validated by quantitative reverse transcription PCR (RT-qPCR). RNA was extracted from each sample using the RNeasy Mini Kit (Qiagen) with optional on-column DNase I digestion. The purity and concentration of the extracted RNA was analyzed using a NanoDrop ND-1000 UV-Vis Spectrophotometer. 1 μg of RNA from each sample was reverse transcribed into cDNA using the High Capacity cDNA Reverse Transcription Kit (Applied Biosystems) according to the manufactures protocol. RT-qPCR was conducted in a 96-well plate format using the Applied Biosystems StepOnePlus Real-Time PCR System. All samples were run in triplicate and each 10 μl reaction contained 2 μL of cDNA (40-fold diluted), 5 μL of 2x Power SYBR™ Green PCR Master Mix (ThermoFisher Scientific), 1 μL each of forward and reverse primers, and 1 μL molecular grade water (Lonza). Each primer pair was tested using a 5-point serial dilution to ensure an efficiency between 90-100% and meltcurve analysis was performed after each run to check for the presence of a single PCR product. The 2^-ΔΔCt^ method was used to determine relative gene expression with elongation factor-1α (EF-1α) serving as the housekeeping gene for normalization [51]. The list of primers used, and their amplification efficiencies are listed in S1 Table.

## Results

### Infection of resistant and susceptible fish stocks

The exposure dose for treatment and control groups (calculated by qPCR) was 7.9 x 10^3^ and 0 actinospores respectively (extrapolated from 1 actinospore standard). Water samples taken at 24 hours were negative, indicating that the spores present successfully attached to the fish. Susceptible fish exposed to *C. shasta* exhibited their first clinical sign of infection at 12 dpe when they stopped responding to feed. At 21 dpe, their intestines were grossly enlarged, inflamed, and bloody, with mature *C. shasta* myxospores visible in swabs of the posterior intestine. Histology revealed a progressive breakdown of the intestinal structure in these fish (Fig 2A,C). By 14 dpe, chronic inflammation could be observed throughout the intestinal submucosa (Fig 2B) and by 21 dpe, all tissue layers were inflamed and sloughing of necrotic mucosal tissue was evident (Fig 2C). No physiological changes were observed by histology in resistant fish (Fig 2D,E).

**Fig 2.**
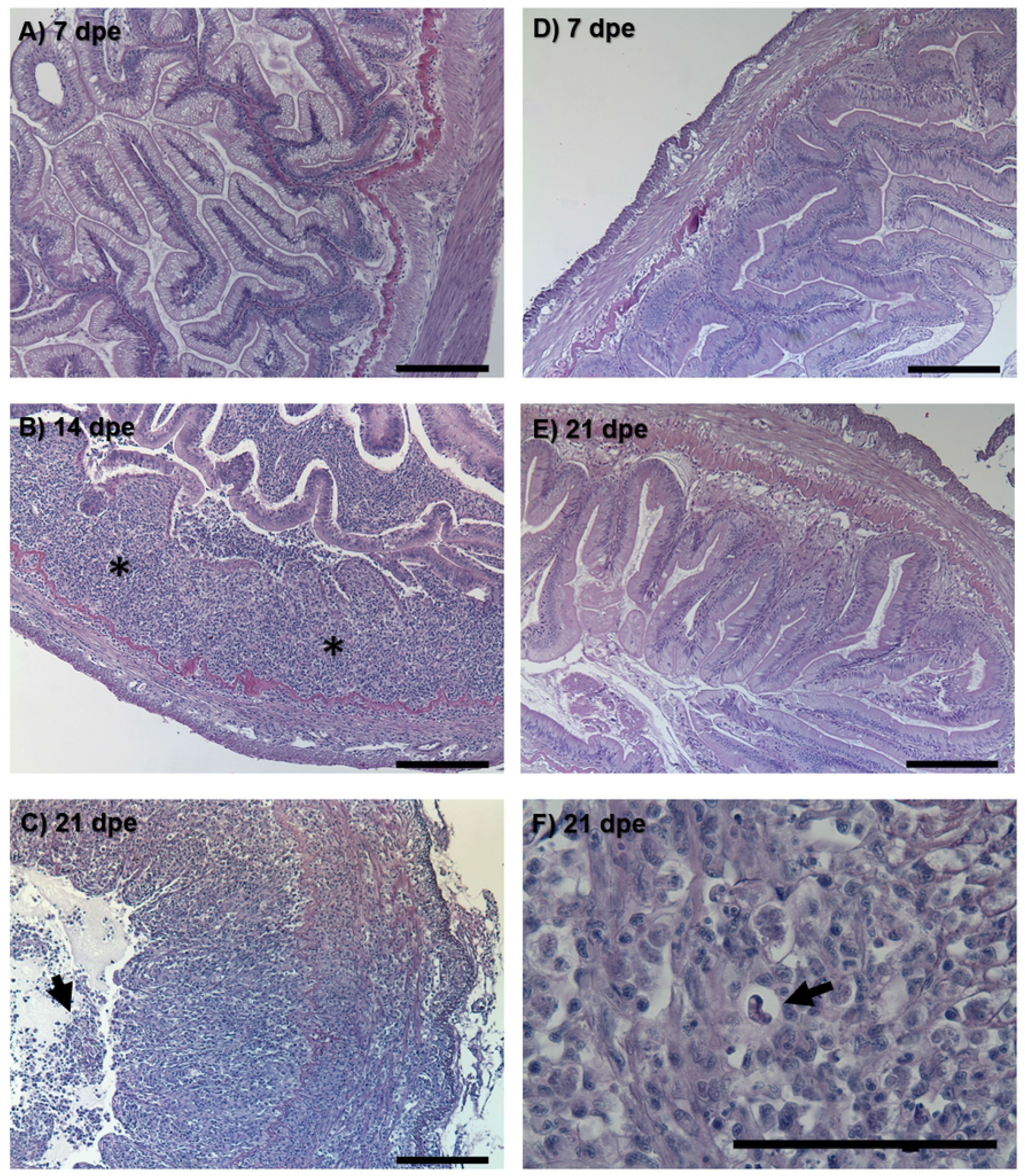
Histological sections of resistant and susceptible steelhead intestine after exposure to *Ceratonova shasta*. Susceptible fish intestine at (A) 7 days post exposure (dpe), (B) 14 dpe showing chronic inflammation (asterisks) throughout the submucosa, and (C) 21 dpe with inflammation present in all tissue layers and sloughing of necrotic epithelia (arrow). Resistant fish intestine at (D) 7 dpe and (E) 21 dpe. Mature *C. shasta* myxospore (arrow) in the intestine of susceptible fish at 21 dpe (F). Bars = 100 μm.

### qPCR quantification of parasite burden

*C. shasta* was not detected by qPCR in the gills at 1 dpe in either the resistant or susceptible fish but was detected in the intestine at 7 dpe in both phenotypes. The infection prevalence among resistant fish remained low throughout the sampling period, with less than half the fish at any timepoint having detectable levels of *C. shasta* in their intestine, and the Cq values of those fish also remained low (31.6 ± 2.2). In contrast, all susceptible fish tested from 7 dpe onwards were positive and had exponentially increasing parasite loads, with Cq values increasing from 24.8 ± 0.8 at 7 dpe to 12.6 ± 0.8 at 21 dpe (Fig 3). No control fish or exposed resistant fish exhibited clinical signs of infection, and randomly selected control fish were negative by qPCR.

**Fig 3.**
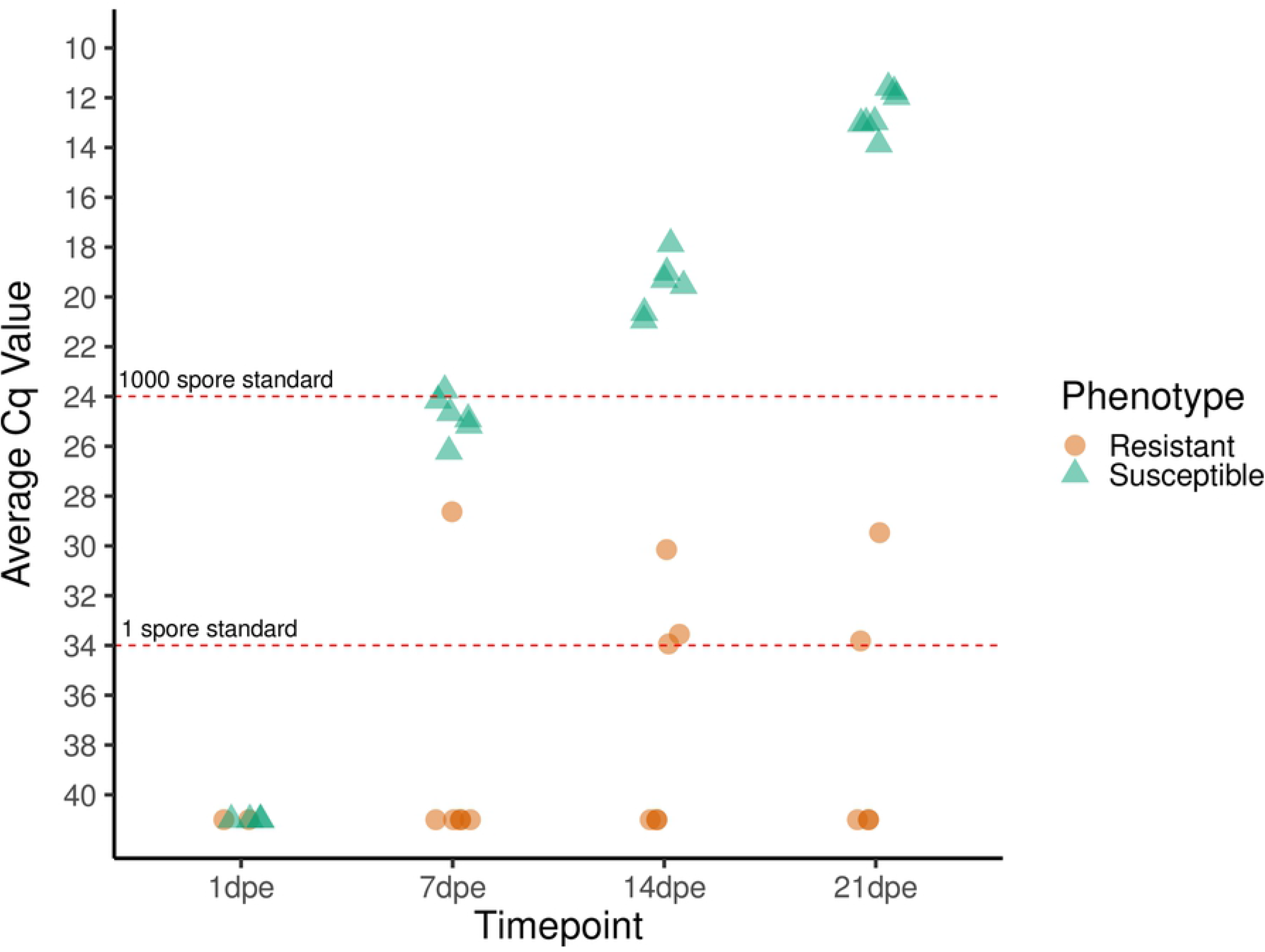
Relative quantity of *Ceratonova shasta* DNA present in the gills (1 dpe) and intestine (7, 14, and 21 dpe) of infected steelhead (*Oncorhynchus mykiss*). Each symbol represents the average quantitative cycle (Cq) of 100 ng of DNA extracted from the whole tissue (gills or intestine) of one fish that was assayed in triplicate by qPCR. Six fish of each phenotype were assayed at each timepoint. Fish that tested negative were assigned a nominal Cq value of 41. Dashed red lines indicate the average Cq values obtained from 1 and 1000 actinospore standards.

### Sequencing

A total of 1.55 x 10^9^ reads were generated from the sequencing of samples from resistant and susceptible fish at 1 and 7 dpe, with an average of 3.22 x 10^7^ (SD ± 4.04 x 10^6^) reads per sample (Table 1). 87.6% of reads could be mapped to the rainbow trout reference genome and 74.8% could be uniquely mapped to specific loci.

**Table 1.** Summary of sequencing results from gill (1 dpe) and intestine (7 dpe) of both resistant and susceptible fish.

| Sequenced Reads |  |
| --- | --- |
| Total | 1,545,135,474 |
| Removed | 518,329 (0.000335%) |
| Mapped | 1,354,217,365 (87.6%) |
| Uniquely Mapped | 1,156,186,486 (74.8%) |
| Average reads per sample | 32,190,322 |

7.80 x 10^8^ reads were generated during the sequencing of samples from susceptible fish at 14 and 21 dpe, with an average of 6.33 x 10^7^ (SD ± 6.00 x 10^6^) reads per sample. The number of reads from exposed susceptible fish that could be mapped to the reference genome decreased to 83.2% at 14 dpe and 42.0% at 21 dpe, reflecting an increase in the amount of parasite RNA present (Table 2).

**Table 2.** Percentage of sequencing reads that mapped to the reference genome at each timepoint.

|  | % of reads mapped |  |  |  |
| --- | --- | --- | --- | --- |
|  | 1 dpe | 7 dpe | 14 dpe | 21 dpe |
| Susceptible - Exposed | 87.5 | 88.0 | 83.2 | 42.0 |
| Resistant - Exposed | 87.7 | 87.4 | - | - |
| Susceptible - Control | 87.3 | 87.8 | 87.2 | 87.9 |
| Resistant - Control | 87.9 | 87.7 | - | - |

### Gills 1 dpe - resistant and susceptible fish - differential gene expression and GO enrichment

The expression of 39,571 genes was detected from sequenced gill transcripts. DEGs responding to *C. shasta* infection were identified by comparing exposed resistant and susceptible fish to their respective controls. This identified 463 DEGs in susceptible fish and 244 in resistant fish, 66 of which were differentially expressed in both phenotypes (Fig 4).

**Fig 4.**
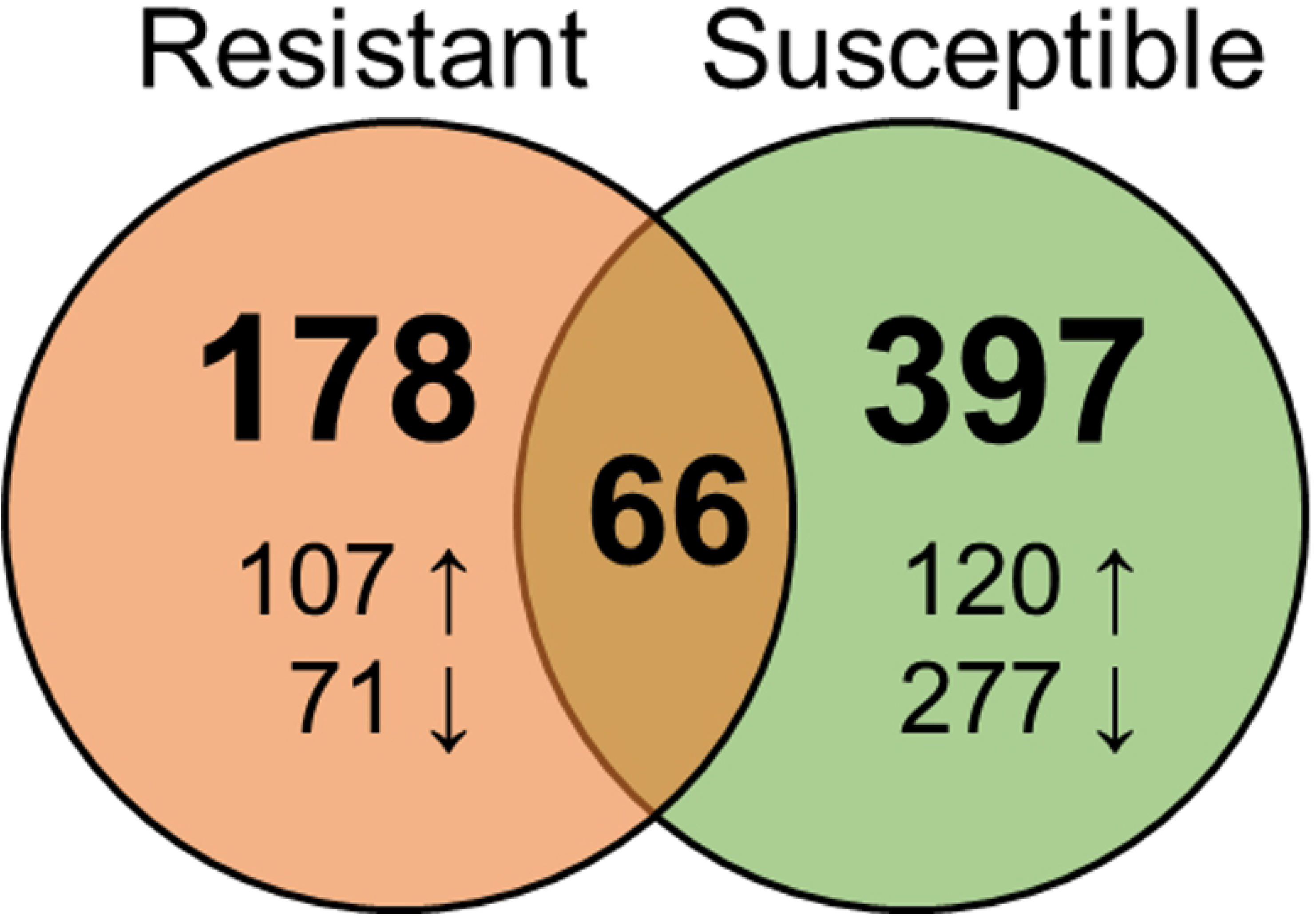
Venn Diagram showing the number of genes differentially expressed in response to *Ceratonova shasta* infection in the gills of resistant and susceptible steelhead at 1 day post exposure. Arrows indicate upregulation vs downregulation.

GO enrichment was conducted to gain insight into the biological processes, molecular functions, and cellular location of the DEGs. In susceptible fish, no specific enrichment was found among the upregulated genes and two GO terms were over-represented among genes upregulated in resistant fish (*carbon dioxide transport* and *one-carbon compound transport*). Among the downregulated genes, resistant fish had 156 enriched GO terms, and susceptible fish had 51. ClueGo analysis revealed that genes involved in the innate immune response, interferon-gamma mediated signaling pathway, response to cytokine, and response to biotic stimulus were over-represented among the downregulated genes for both resistant and susceptible fish (Fig 5). Many of the downregulated immune genes were shared by both phenotypes (Table 3), including interferon gamma 2, Interferon-induced protein 44, and several C-C motif chemokines.

**Fig 5.**
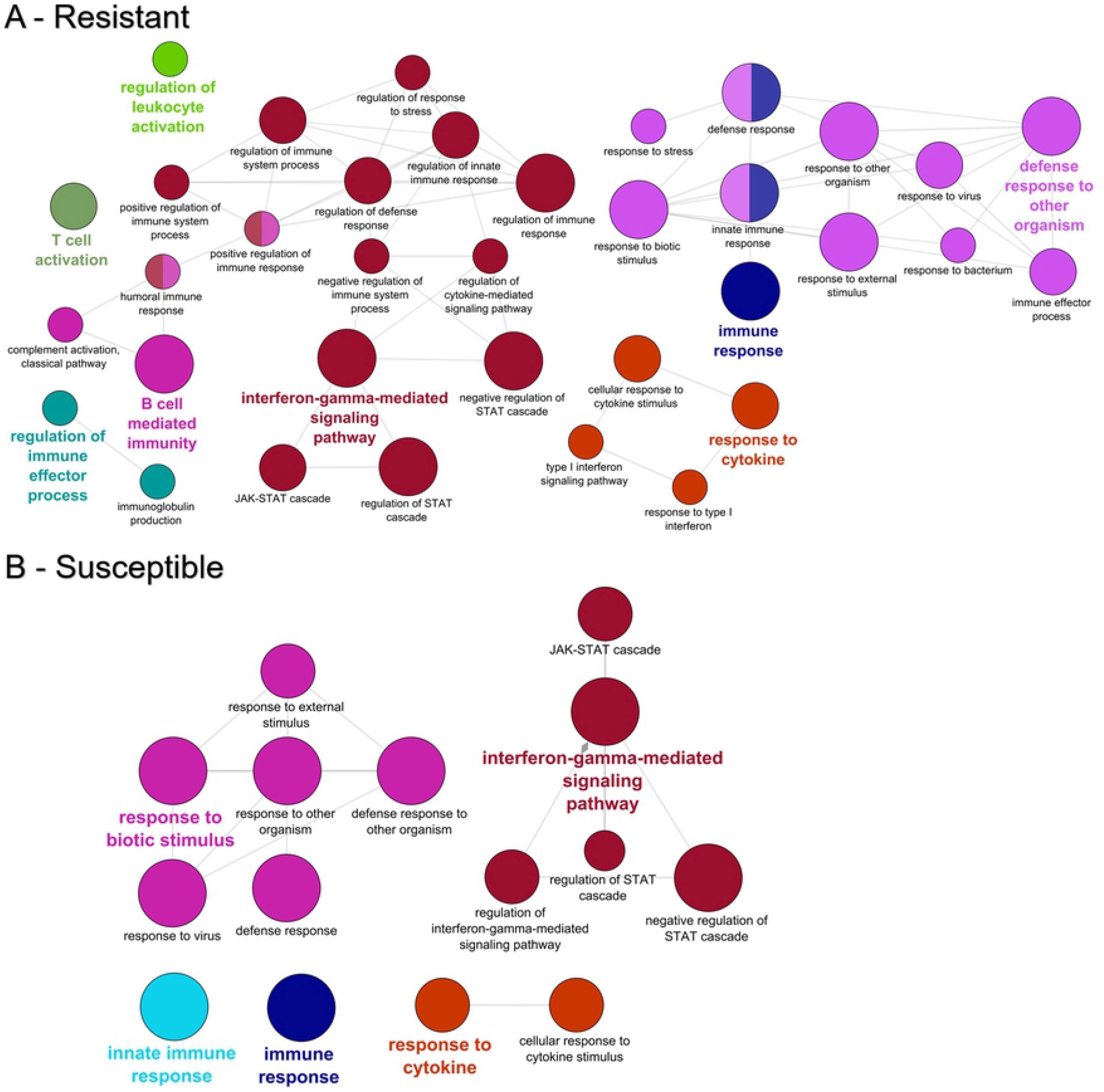
GO enrichment among the genes downregulated in the gills of resistant (A) and susceptible (B) steelhead at 1 day post exposure to *Ceratonova shasta*. Enriched gene ontology (GO) terms were grouped into functionally related nodes using the Cytoscope plugin ClueGO. Nodes are colored and grouped according to a related function and labelled by the most significant term of the group. Node size corresponds to the FDR-adjusted p-value of each GO term and is specific to each graph.

**Table 3.**
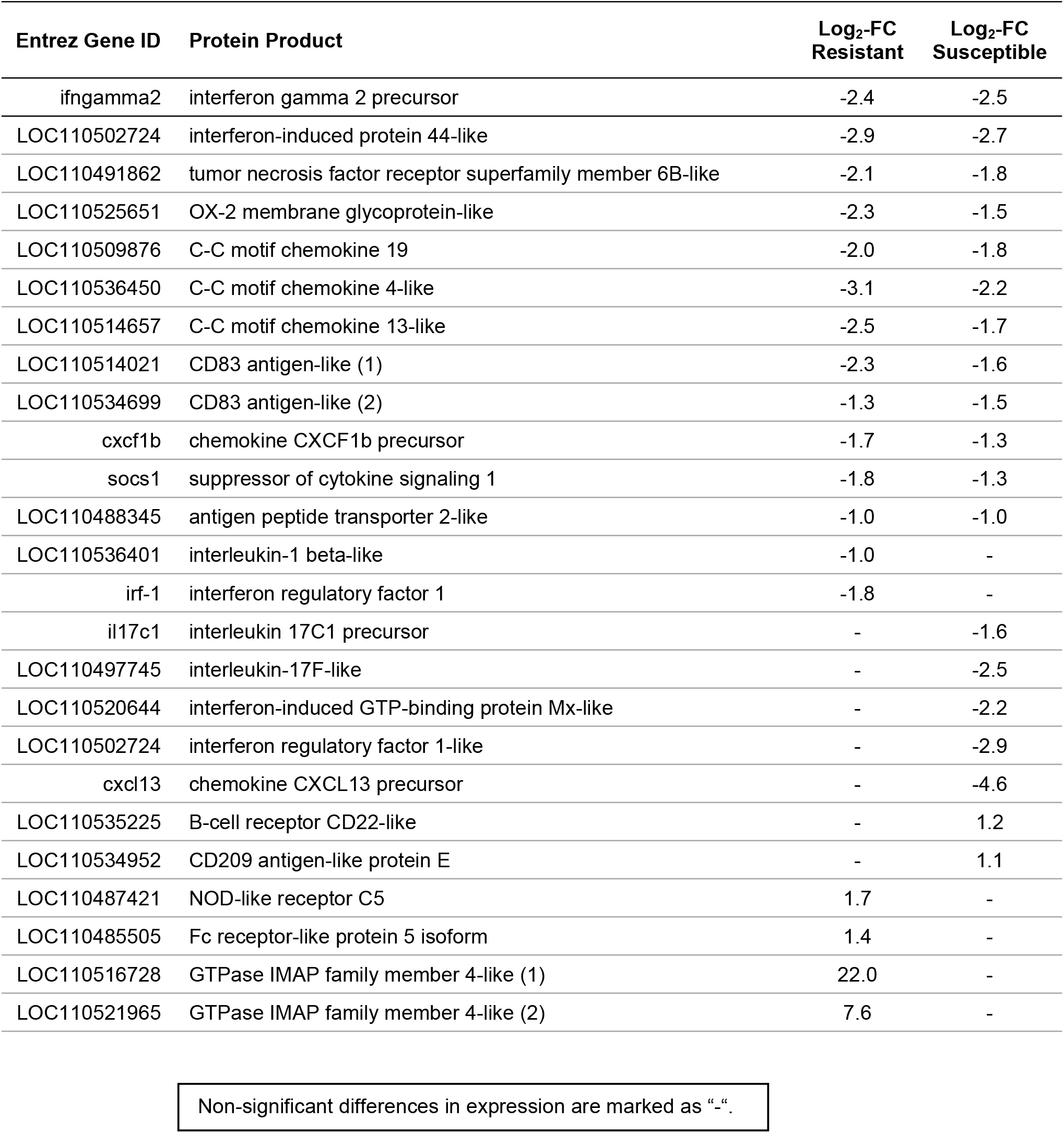
Select immune genes that were differentially expressed in the gills of resistant and susceptible steelhead at 1 day post exposure to *Ceratonova shasta.* Non-significant differences in expression are marked as Non-significant differences in expression are marked as “-”.

While most immune related DEGs were downregulated in both phenotypes, the two most highly upregulated genes in resistant fish were homologs of GTPase IMAP family member 4-like at 22.0 and 7.6 log_2_-FC, respectively. GIMAPs (GTPase of the immunity associated protein family) are a relatively recently described family of small GTPases that are conserved among vertebrates and are associated with T-lymphocyte development and activation [52]. Two immune receptors were also upregulated in resistant fish: NLRC 5 and Fc receptor-like protein 5. In susceptible fish, only two immune genes were upregulated: B-cell receptor CD22-like and CD209 antigen-like protein E.

### Intestine 7 dpe - resistant and susceptible fish - differential expression and GO enrichment

37,978 genes were identified in the intestine at 7 dpe. As for gills, DEGs were identified by comparing exposed fish to their unexposed controls. In contrast to the large number of DEGs in the gills at 1 dpe, only 16 DEGs were identified in resistant fish, 4 in susceptible fish, and no DEGs overlapped between them (Table 4). No GO enrichment was conducted due to the small number of DEGs.

**Table 4.**
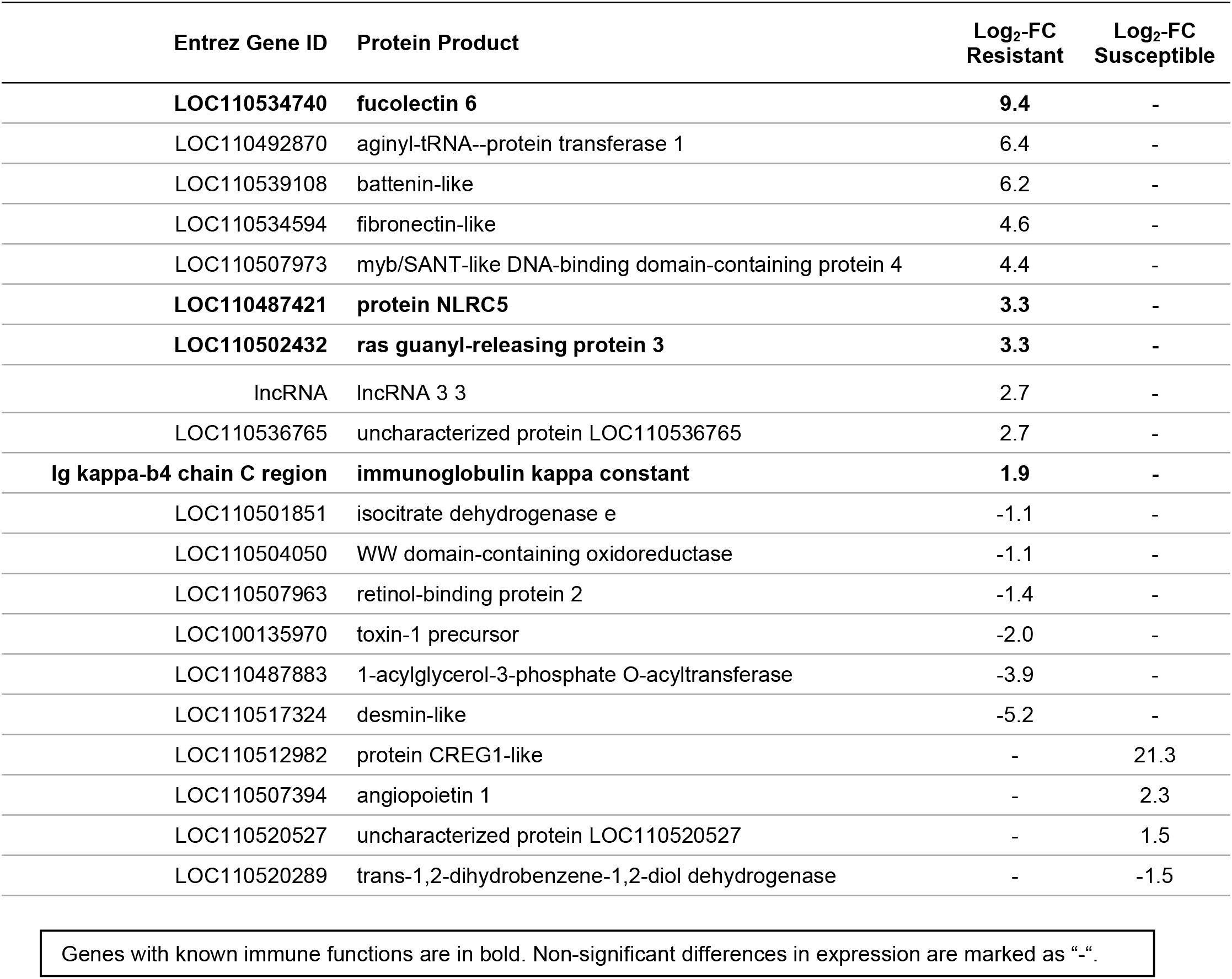
Genes that were differentially expressed in the intestine of resistant and susceptible steelhead 7 days post exposure to *Ceratonova shasta.* Genes with known immune functions are in bold. Non significant differences in expression are marked asGenes with known immune functions are in bold. Non-significant differences in expression are marked as “-”.

Among the DEGs in resistant fish that have known functions, four immune genes were upregulated, including two innate immune receptors: Fucolectin 6, an F-type lectin that binds fucose, and NLRC 5, which was also upregulated in the gills of resistant fish at 1 dpe. Two immune genes involved in B cell responses were also upregulated: Ras guanyl-released protein 3, involved in B cell activation [53], and immunoglobulin kappa constant. Fibronectin-like, an extracellular matrix protein, and battenin-like were also significantly upregulated. Battenin, also called CLN3, is a highly conserved multi-pass membrane protein that localizes to the lysosome and other vesicular compartments, but the function of which remains unknown [54]. The most downregulated gene in resistant fish was desmin-like protein, a muscle specific intermediate filament.

In susceptible fish, the cell-growth inhibitor protein CREG1 was the most highly upregulated transcript, followed by the vascular growth factor angiopoietin-1-like.

### Comparison of resistant and susceptible controls

To identify any genes involved in resistance to *C. shasta* that might be constitutively expressed in resistant fish, we conducted a differential gene expression analysis comparing the uninfected controls for both phenotypes. This yielded 1400 DEGs in the gills, and 307 in the intestine. 38 DEGs were present in both tissues and upregulated in resistant fish relative to susceptible fish (S2 Table). Among them were six genes associated with immune system functions: two homologs of NLRC 5 (not the same one upregulated in response to *C. shasta* infection), GTPase IMAP family member 7-like, complement C1q-like protein 2, TGF-beta receptor type-2-like, and perforin-1-like.

### Intestine - susceptible fish - 14 and 21 dpe - differential gene expression and GO enrichment

The transcriptomic response of susceptible fish was followed through later timepoints to determine how these fish reacted as the parasite continued to proliferate. Sequencing of infected fish and their time-matched controls identified 36,957 and 36,346 gene transcripts at 14 and 21 dpe, respectively. Comparison to the intestine of uninfected susceptible fish revealed 5,656 DEGs at 14 dpe and 12,061 DEGs at 21 dpe, 3,708 of which were differentially expressed at both timepoints (Fig 6).

**Fig 6.**
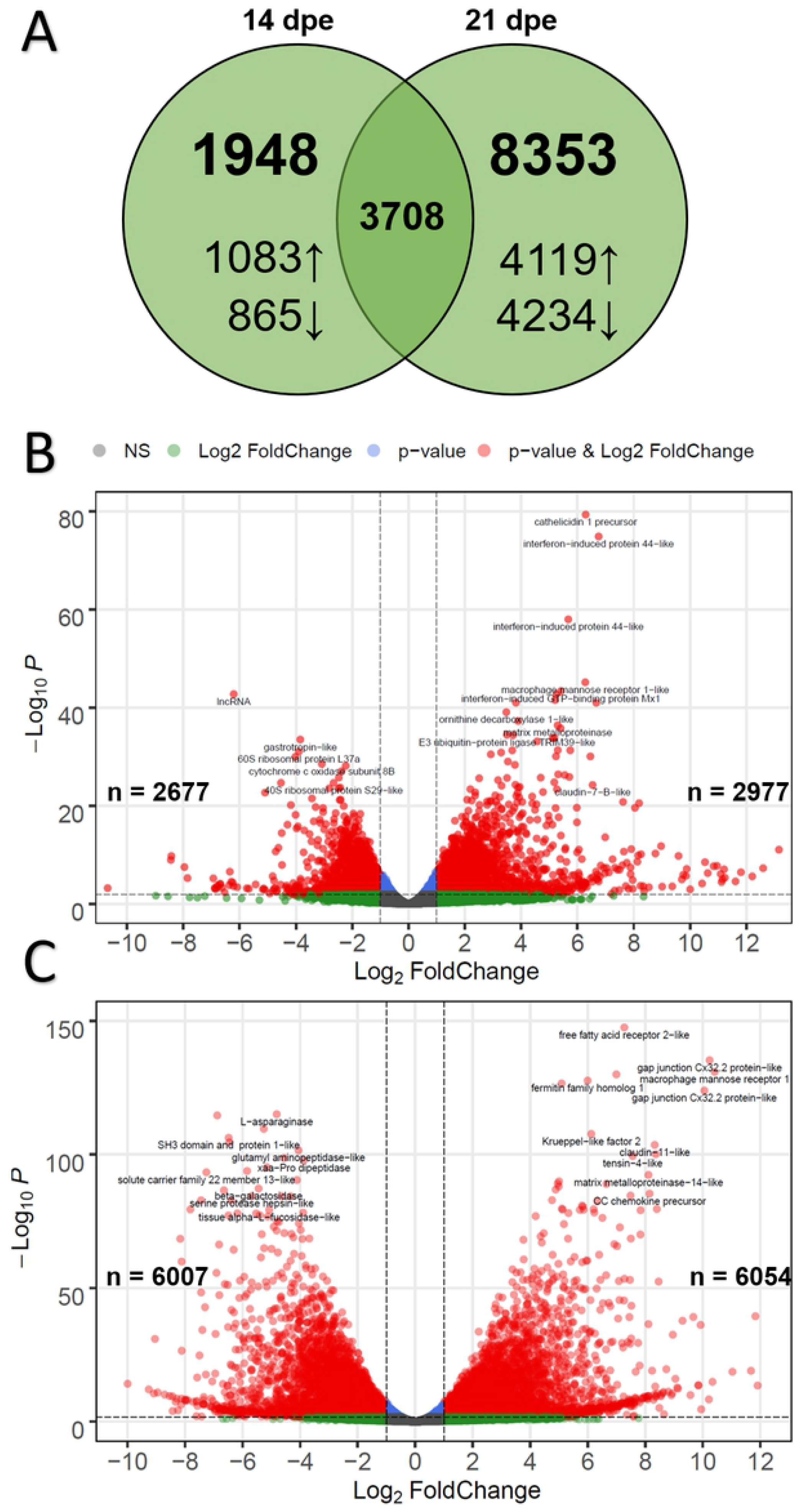
Differential expression results for susceptible fish at 14- and 21-days post exposure (dpe) to *Ceratonova shasta*. A) Venn diagram indicating the number of differentially expressed genes overlapping at 14- and 21 dpe. Arrows indicate up-vs. downregulation. B) Volcano plot of differential gene expression for susceptible fish at 14 dpe. Each dot represents the average value of one gene across three biological replicates. Red indicates the gene was significant at the FDR-adjusted p-value and Log_2_-Foldchange threshold, blue is significantly only by p-value, green only by Log_2_-Foldchange, and gray were not significant by either metric. B) Same as (A), but for susceptible fish at 21 dpe.

GO enrichment analysis of the 2,977 upregulated genes at 14 dpe indicated 631 overrepresented GO terms, primarily immune related. ClueGO analysis clustered these into networks revolving around GO terms for interferon-gamma-mediated signaling pathway, regulation of defense response, positive regulation of response to external stimulus, immune response, and innate immune response (Fig 7A). The same analysis for the 2,677 downregulated genes at 14 dpe yielded 196 GO terms, which clustered into networks based on terms for oxidation-reduction process, mitochondrion organization, translation, and lipid catabolic process (Fig 7B).

**Fig 7.**
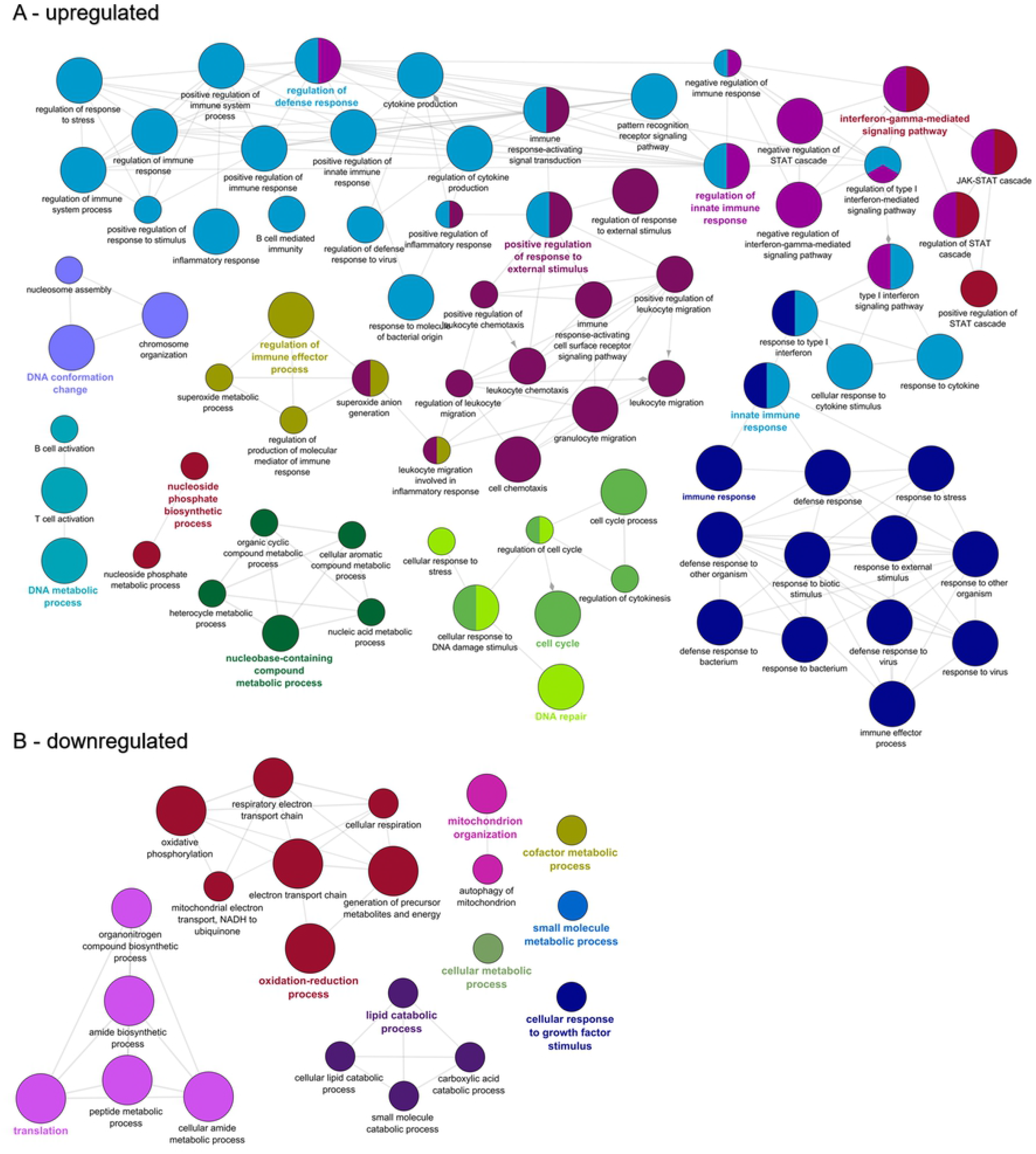
Functional enrichment of biological processes among the genes differentially expressed in the intestine of susceptible fish at 14 days post exposure to *Ceratonova shasta*. Enriched gene ontology (GO) terms were grouped into functionally related nodes using the Cytoscope plugin ClueGO. Nodes are colored and grouped according to a related function and labelled by the most significant term of the group. Node size corresponds to the FDR-adjusted p-value of each GO term and is specific to each graph. The analysis was conducted separately on upregulated (A) and downregulated (B) genes.

At 21 dpe, the 6,054 upregulated genes contained 452 over-represented GO terms which primarily clustered into networks revolving around immune system processes such as immune response-activating signal transduction, positive regulation of immune system process, immune response-activating cell surface receptor signaling pathway, and regulation of immune response (Fig 8A). In addition to these immune system pathways, cell adhesion pathways came to the forefront, including cell-matrix adhesion, cytoskeleton organization, integrin-mediated signaling pathway, and positive regulation of cell adhesion. The 6,007 downregulated genes were enriched for 152 GO terms that clustered into networks for lipid catabolic process, oxidation-reduction process, lipid metabolic process, and cofactor metabolic process (Fig 8B).

**Fig 8.**
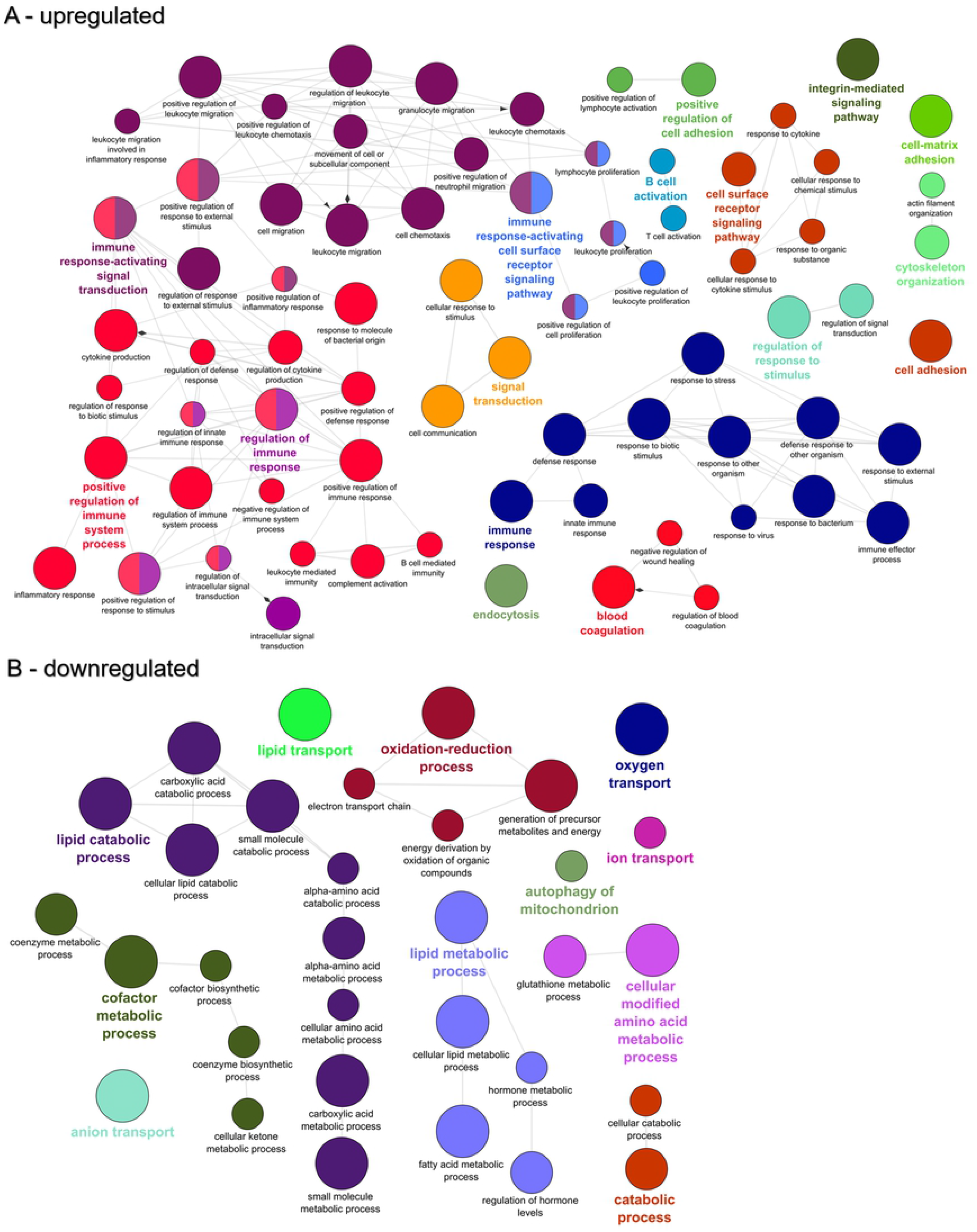
Functional enrichment of biological processes among the genes differentially expressed in the intestine of susceptible fish at 21 days post exposure to *Ceratonova shasta*. Enriched gene ontology (GO) terms were grouped into functionally related nodes using the Cytoscope plugin ClueGO. Nodes are colored and grouped according to a related function and labelled by the most significant term of the group. Node size corresponds to the FDR-adjusted p-value of each GO term and is specific to each graph. The analysis was conducted separately on upregulated (A) and downregulated (B) genes.

### Key genes expressed in response to *C. shasta* infection in susceptible fish

Due to the large number of DEGs detected, only a subset of key genes identified in our analysis are presented in Table 5 and described below. The complete list of differential gene expression results and GO enrichment can be found in S2 Table.

**Table 5.** Select immune genes that were differentially expressed in the intestine of susceptible steelhead at 14- and 21-days post exposure (dpe) to *Ceratonova shasta.*

| Entrez Gene ID | Protein Product | Log<sub>2</sub>-FC<br>14 dpe | Log<sub>2</sub>-FC<br>21 dpe |
| --- | --- | --- | --- |
| **Cytokines** |  |  |  |
| LOC100136024 | interleukin-1 beta | 3.3 | - |
| LOC110536401 | interleukin-1 beta-like | 9.0 | 5.3 |
| il-6 | interleukin-6 precursor | - | 5.9 |
| LOC110496949 | interleukin-6-like | 7.5 | 10.3 |
| il-8 | putative CXCL8/interleukin-8 | 2.6 | 5.3 |
| LOC110531606 | CXCL8/interleukin-8-like | 1.9 | 5.9 |
| tnf | tumor necrosis factor | - | - |
| mif | macrophage migration inhibitory factor | -2.0 | -2.7 |
| LOC110488642 | macrophage migration inhibitory factor-like | -2.2 | -2.2 |
| csf-3 | granulocyte colony-stimulating factor precursor | 4.4 | - |
| csf3r | granulocyte colony-stimulating factor receptor | 1.6 | 3.0 |
| **Effector enzymes** |  |  |  |
| LOC110536463 | granzyme A-like | 5.2 | 2.7 |
| LOC110520655 | granzyme B-like | 3.9 | 5.0 |
| LOC110524258 | granzyme-like protein 2 | - | 2.9 |
| LOC110531658 | perforin-1-like | 1.7 | 1.4 |
| LOC110536422 | perforin-1-like | -1.7 | -1.2 |
| LOC110538116 | perforin-1-like | 3.9 | 7.4 |
| LOC110500520 | perforin-1-like | - | 5.0 |
| LOC100136187 | cathelicidin antimicrobial peptide | 5.8 | 8.3 |
| LOC100136204 | cathelicidin 1 precursor | 6.3 | 7.5 |
| LOC110523157 | lysozyme C II | - | 2.5 |
| LOC110485102 | lysozyme g-like | - | 2.0 |
| **Macrophages** |  |  |  |
| LOC110498289 | arginase-2, mitochondrial-like | - | 4.1 |
| LOC110506002 | arginase-2, mitochondrial-like | 4.1 | 4.0 |
| nos2 | nitric oxide synthase, inducible | - | - |
| LOC110507147 | nitric oxide synthase, inducible (Fragment)-like | 4.7 | - |
| LOC110536912 | macrophage mannose receptor 1-like | -1.9 | -6.7 |
| LOC110500089 | macrophage mannose receptor 1-like | - | -3.9 |
| LOC110508265 | macrophage mannose receptor 1-like | 6.3 | 10.4 |
| LOC110508267 | macrophage mannose receptor 1-like | 5.5 | 7.8 |
| LOC110516203 | macrophage mannose receptor 1-like | 5.0 | 8.2 |
| **T<sub>H1</sub> response** |  |  |  |
| ifng | interferon gamma | - | 6.6 |
| ifngamma2 | interferon gamma 2 | 5.4 | 4.7 |
| ifngr1 | interferon gamma receptor 1 | 4.2 | 3.5 |
| ifngr1 | interferon gamma receptor alpha chain precursor | - | 1.6 |
| irf-8 | interferon regulatory factor 8-like | 3.7 | 2.5 |
| il12b | interleukin-12 beta chain precursor | 2.1 | 2.2 |
| LOC110537792 | interleukin-12 subunit beta-like | - | -4.0 |
| LOC110524480 | interleukin-12 receptor subunit beta-2-like | 1.1 | -1.3 |
| LOC110524481 | interleukin-12 receptor subunit beta-2-like | 1.5 | 2.9 |
| LOC110511354 | interleukin-18 receptor accessory protein-like | - | 5.0 |
| tbx21 | T-bet | 2.4 | 4.3 |
| stat1-1 | signal transducer/activator of transcription 1 | 2.1 | 1.3 |
| LOC110520020 | signal transducer and activator of transcription 1-alpha/beta-like | 3.8 | 3.0 |
| LOC110501544 | signal transducer and activator of transcription 1-alpha/beta-like | 2.5 | 3.2 |
| **T<sub>H2</sub> response** |  |  |  |
| il4/13a | interleukin-4/13A precursor | 5.4 | 5.0 |
| LOC110489171 | interleukin-4/13b1 precursor | 6.1 | 5.0 |
| LOC110504551 | interleukin-4/13b2 precursor | 7.9 | 7.7 |
| il17c1 | interleukin-17C1 precursor | - | -6.8 |
| LOC110492428 | interleukin-17 receptor C-like | - | -2.7 |
| socs3 | suppressor of cytokine signaling 3 | 4.1 | 4.1 |
| LOC110512513 | suppressor of cytokine signaling 3-like | 3.9 | 3.9 |
| LOC110500122 | transcription factor GATA-3-like | - | 2.2 |
| **T<sub>H17</sub> response** |  |  |  |
| il-17a | interleukin-17A precursor | -8.4 | -6.7 |
| LOC110504334 | interleukin-17A-like | - | -5.4 |
| LOC110529296 | interleukin-17A-like | 1.4 | - |
| il-17d | interleukin-17 isoform D precursor | -2.1 | - |
| LOC110497745 | interleukin-17F-like | - | -2.0 |
| LOC110505720 | interleukin-17F-like | - | -8.3 |
| il17rd | interleukin-17 receptor D | - | -1.2 |
| LOC110492331 | interleukin-17 receptor D-like | 1.4 | 2.0 |
| il17r | interleukin-17 receptor precursor | - | -2.4 |
| il-22 | interleukin-22 precursor | - | -3.1 |
| LOC110524663 | interferon regulatory factor 4-like | 1.3 | 1.4 |
| LOC110538194 | signal transducer and activator of transcription 3 | - | -1.2 |
| LOC110520784 | nuclear receptor ROR-gamma-like | - | -2.8 |
| LOC110535950 | nuclear receptor ROR-gamma-like | - | -2.4 |
| **T<sub>reg</sub> response** |  |  |  |
| il10 | interleukin-10 precursor | 6.3 | 8.1 |
| il10b | interleukin-10b protein precursor | 4.8 | 6.0 |
| LOC100136774 | transforming growth factor beta-1 | 1.4 | 1.4 |
| LOC110534057 | transforming growth factor beta-1-like | 1.7 | 3.9 |
| tgfb1i1 | transforming growth factor beta-1-induced transcript 1 protein | - | 2.4 |
| foxp3-1 | forkhead box P3-1 protein | - | -1.9 |
| foxp3-2 | forkhead box P3-2 protein | - | -2.3 |
| **B cell response** |  |  |  |
| LOC110522002 | Blimp-1/PR domain zinc finger protein 1-like | 4.9 | 7.5 |
| LOC110496128 | Blimp-1/PR domain zinc finger protein 1-like | 3.3 | 4.2 |
| LOC110485501 | B-cell receptor CD22-like | 2.5 | 5.2 |
| LOC110538709 | immunoglobulin heavy variable 1-69-2-like | 5.7 | 8.0 |
| LOC110490545 | Ig kappa chain V region K29-213-like | 2.3 | 2.4 |
| LOC110535024 | immunoglobulin kappa light chain-like | 2.1 | 2.1 |

Non-significant differences in expression are marked as “-”.

### Cytokines

The pro-inflammatory cytokine interleukin-1 beta (IL-1β) was highly upregulated at 14- and 21 dpe. IL-1β is a chemoattractant for leukocytes in fish and modulates the expression of other chemokines including CXCL8/interleukin-8 [55], which was also upregulated at both timepoints. Curiously, the pro-inflammatory cytokine TNA-α was not differentially expressed at either timepoint despite the upregulation of other pro-inflammatory cytokines, including IL-1β which stimulates the production of TNA-α. Macrophage migration inhibitory factor (MIF) is a pro-inflammatory cytokine that acts as a mediator of both innate and acquired immunity. It is implicated in resistance to bacterial pathogens and is released from macrophages after stimulation with LPS. Mice that lack MIF are more susceptible to leishmaniasis and cysticercosis and *in vivo* administration of recombinant MIF reduced the severity of *Leishmania major* pathogenesis in mice [56]. We observed downregulation of two MIF homologs at both 14 and 21 dpe.

### Effector enzymes

We detected low to high upregulation of several granzyme and perforin transcripts at 14- and 21 dpe. Cytotoxic T lymphocytes (CTLs) release these proteins in secretory granules to induce apoptosis of infected or damaged cells. The antimicrobial peptide cathelicidin was highly upregulated at both timepoints, while lysozyme was upregulated only at 21 dpe.

### Macrophage activation and polarization

Macrophages at the site of inflammation polarize into M1 or M2 phenotypes. M1 polarization is associated with the T_H1_ response and the presence IFN-γ and induces macrophages to express the enzyme nitric oxide synthase (NOS) leading to the production of reactive nitrogen species for pathogen clearance. M2 polarization is driven by the T_H2_ response and the presence IL-4/13. M2 macrophages are associated with wound healing and the expression of the arginase enzyme. We observed upregulation of NOS at 14 dpe but not at 21 dpe. The opposite was true for arginase, which was only upregulated at 21 dpe.

Macrophage mannose receptor 1 (MCR1) is a transmembrane glycoprotein belonging to the C-type lectin family. In addition to scavenging certain hormones and glycoproteins, it also recognizes a variety of pathogens including influenza virus, *Yersinia pestis*, and *Leishmania* species [57]. Ten homologs of MCR1 were differentially expressed at 14- or 21 dpe and were among the most highly induced immune genes at 21 dpe.

### GTPase IMAP family members

A total of 15 GIMAP proteins were upregulated at 14 dpe, including two homologs of GTPase IMAP family member 4-like which were the two most highly upregulated immune genes at this timepoint (10.4 and 9.0 log_2_-FC). The same two homologs were also the most highly upregulated genes (22.0 and 7.6 log_2_-FC) in the gills of resistant fish at 1 dpe. However, they were not differentially expressed in susceptible fish at 21 dpe. At 21 dpe, only 5 GIMAPs proteins were upregulated, with GTPase IMAP family member 7-like having the highest increase in expression (4.1 log_2_-FC).

### Activated T-cells

CD4^+^ T helper cells (T_H_ cells) are an important wing of the adaptive immune response that differentiate into one of several effector subsets (T_H1_, T_H2_, T_H17_, and T_reg_) based on the cytokine signals they receive. These effector cells, in turn, secrete their own district profile of cytokines that help orchestrate the immune response. Among the genes differentially expressed in response to *C. shasta* infection, signature genes for each subset were identified to provide insight into the T cell response (Table 6).

Interferon gamma (IFN-γ), the signature T_H1_ cytokine, was highly upregulated at both 14- and 21 dpe along with its cognate receptor and T-bet, the master transcriptional regulator of T_H1_ differentiation. Only one gene related to interleukin-12, the primary driver of T_H1_ differentiation, was upregulated at 14 dpe (interleukin-12 subunit beta-like, 2.3 Log_2_-FC). The gene was similarly upregulated at 21 dpe, along with interleukin-12 alpha and beta chains.

Interleukin-4/13 is the primary cytokine produced by T_H2_ cells and drives alternative macrophage activation and type 2 inflammation. Moderate upregulation of interleukin-4/13A precursor was seen at 14- and 21 dpe. The master transcriptional regulator of T_H2_ differentiation, GATA-3, was only upregulated at 21 dpe.

Downregulation of several genes involved in the T_H17_ response was observed at 14- and 21 dpe. Most significant of these was interleukin-17A precursor, and interleukin-17F-like. Two copies of nuclear receptor ROR-gamma, the putative master transcriptional regulator of T_H17_, were downregulated at 21 dpe.

Little evidence of a strong regulatory T cell response was seen at either 14- or 21 dpe. FOXP3, the master transcriptional regulator for T_reg_ cells, was downregulated at 21 dpe. Transforming growth factor beta transcripts were mildly upregulated at both timepoints. Interleukin-10, which is classically associated with T_reg_, was highly upregulated at 14- and 21 dpe, however, it can be produced by numerous different myeloid and lymphoid cells during an infection [58]. This lack of an observable T_reg_ response may be due to the significant upregulation of interleukin-6 seen at both timepoints, as interleukin-6 is known to inhibit T_reg_ conversion in humans and mice [59,60].

### B cell response

Numerous genes involved in the B cell response and production of immunoglobulins were upregulated at 14 dpe, and both the number of genes and the magnitude of the upregulation increased at 21 dpe. Among these were the transcription factor Blimp-1, which is required for the maturation of B cells into Ig-secreting cells, B cell receptor CD22, and several heavy and light chain transcripts.

### Innate immune receptors

Toll-like receptors (TLRs) are innate immune receptors that recognize conserved pathogen-associated molecular patterns. We observed upregulation of six different TLRs at 14- or 21 dpe, including eleven homologs of TLR13. In mice, TLR13 recognizes a conserved bacterial 23S ribosomal RNA sequence, a function that appears to be conserved in teleost fish [61]. Two copies of TLR8, which recognizes viral single-stranded RNA, were upregulated at both timepoints, and one copy of TLR1, which recognizes bacterial lipoprotein, and TLR22. TLR22 is a fish-specific TLR and has been shown to be induced after viral, bacterial, or ectoparasite challenge [62]. TLR3 and TLR7, which recognize viral RNA, were upregulated at 14 dpe. Although they were different homologs than those upregulated in resistant fish, 18 putative NOD-like receptors were upregulated at 14 dpe and 11 at 21 dpe. We also observed substantial upregulation of C-type lectins, with 16 upregulated at 14 dpe and 20 upregulated at 21 dpe.

### Cell adhesion

Genes involved in cell-to-cell contact and the formation of the intestinal barrier were among the most transcriptionally active at both timepoints, with the majority of transcripts being upregulated. At 14 dpe, this included 10 claudins, 19 integrins, 1 fibronectin, 5 fermitin family homologs, 8 gap junction proteins, and 17 cadherins. This continued at 21 dpe with 23 claudins, 42 integrins, 11 fibronectins, 7 fermitin family homologs, 15 gap junction proteins and 36 cadherins. Additionally, in terms of statistical significance, the actin binding protein beta-parvin was the most significant DEG at 21 dpe (padj = 9.97e-232, log_2_-FC = 7.6).

### Validation of DEGs using RT-qPCR

Four immune genes *(IFN-y TNF-α, IL-10, IL-1β)* found to be differentially expressed by RNA-seq were assayed using quantitative reverse transcription PCR (RT-qPCR) to validate the results and confirm the observed downregulation of immune genes. Fold changes from RT-qPCR are compared with those from RNA-seq in Fig 9 and support the results we obtained.

**Fig 9.**
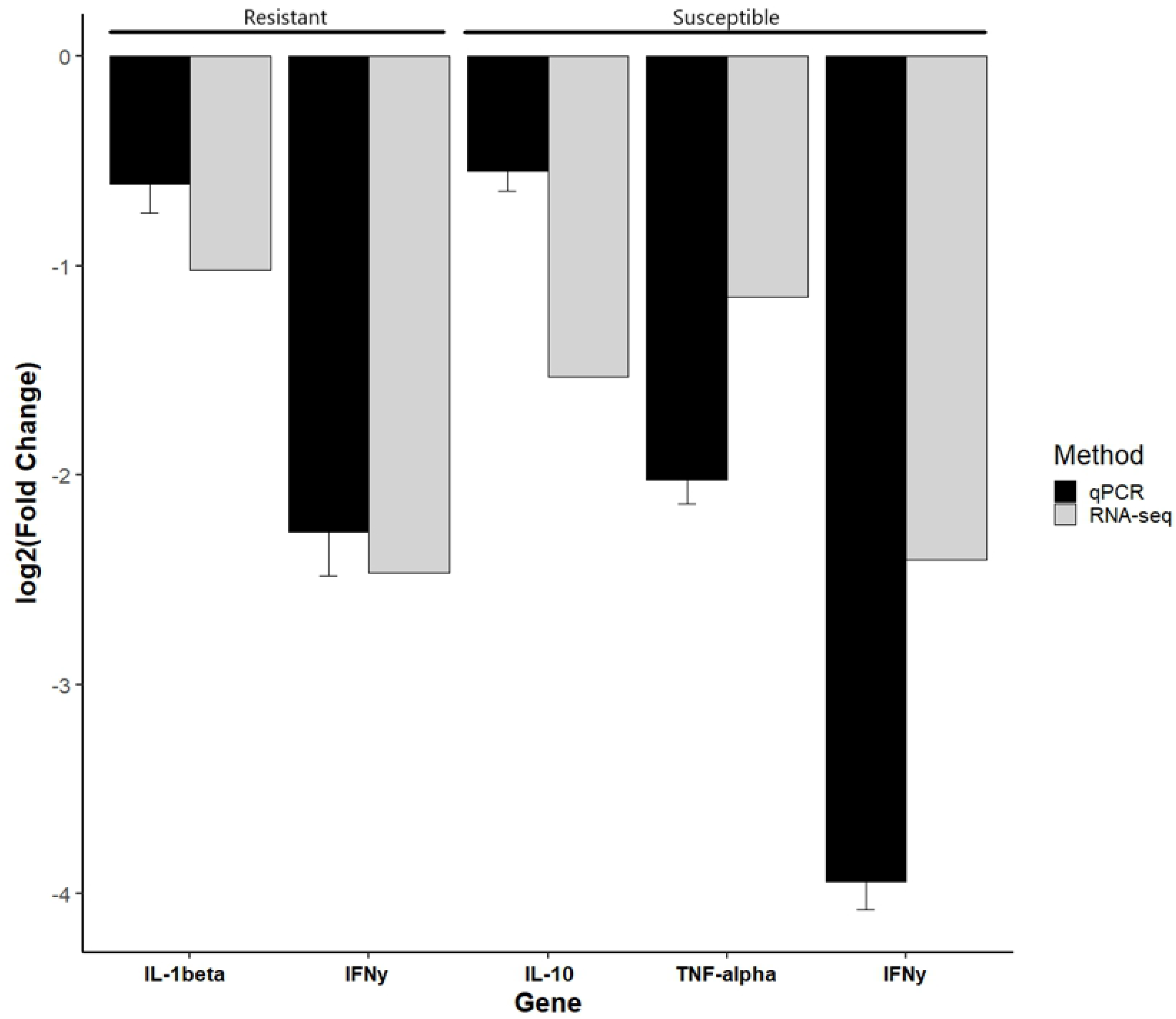
qPCR validation of RNA-seq results. Quantitative reverse transcription PCR (RT-qPCR) validation of four immune genes (*IFN-y TNF-α, IL-10, IL-1β*) found to be significantly differentially expression by RNA-seq at day 7 in the gills. The X-axis shows the gene and phenotype assayed and the Y-axis shows the relative log_2_(Fold Change) between fish exposed to *Ceratonova shasta* and their respective control. Error bars indicate the standard deviation of Cq values between biological replicates.

## Discussion

We used RNA-seq to study the early transcriptomic response of resistant and susceptible steelhead infected with the myxozoan parasite *C. shasta.* Comparative transcriptomics revealed that both phenotypes had a suppression of the interferon gamma signaling pathway in the gills at 1 dpe. The response of the two phenotypes quickly diverges after that. In the intestine at 7 dpe, resistant fish had effectively contained the parasite and several immune genes were upregulated in this tissue. Susceptible fish, on the other hand, had no observable response to parasite proliferation in the intestine at this time. Parasite replication in susceptible fish continued exponentially at 14- and 21 dpe, which coincided with an intense, yet ineffective immune response and the breakdown of the intestinal structure.

### Immunosuppression at the portal of entry (gills)

Given the markedly different resistance of these two fish stocks to *C. shasta* induced pathology, the overall transcriptomic response in the gills was surprisingly similar, with a downregulation of immune genes in both phenotypes. We observed a suppression of the innate immune response, particularly the IFN-γ signaling pathway which is the primary immune pathway activated later in the infection. This may reflect a parasite-induced immunosuppression that aids in initial invasion of the host. Immunosuppression is a well-known method of immune evasion for human parasites [63], and an immunosuppressed state has been observed in other fish-parasite systems, including infections by other myxozoans. A microarray analysis of gilthead sea bream exposed to the myxozoan *Enteromyxum leei* revealed that successfully parasitized fish were characterized by a global downregulation of genes involved in the immune and acute phase response [64]. Studies of rainbow trout infected with the related malacosporean *Tetracapsuloides bryosalmonae*, the causative agent of proliferative kidney disease, revealed suppression of phagocytic activity and oxidative burst [65], and a dysregulated T-helper and B cell response [66,67]. The transcriptomic response of Atlantic salmon affected by amoebic gill disease, caused by a protozoan parasite, is also associated with downregulation of immune genes, including those related to MHC I and IFN-γ [68,69].

### Potential recognition of the parasite by resistant fish

Although the majority of immune genes were downregulated in the gills of resistant fish, two copies of GTPase IMAP family member 4-like were the most highly upregulated genes at this timepoint. Additionally, the immune receptors NLRC5 and Fc receptor-like protein 5 were also upregulated. The upregulation of innate immune receptors, including NLRC 5 which was also upregulated in the intestine of resistant fish, suggests that specific recognition of *C. shasta* may be occurring in these fish. While this may not offer protection at the portal of entry, it may enable a more rapid immune response to the parasite at the intestine, or during its migration there. This would explain why resistant fish had a much lower infection prevalence and intensity in the intestine (Fig 3).

### GIMAPs may mediate resistance to *C. shasta*

As noted above, the two most highly upregulated genes in the gills of resistant fish were two homologs of GTPase IMAP family member 4-like, a protein involved in T-lymphocyte development. Intriguingly, the same two homologs were the most highly upregulated immune genes in the intestine of susceptible fish at 14 dpe. If these genes are involved in mediating resistance to *C. shasta,* then their delayed expression in susceptible fish could explain the delayed immune response observed in these fish. How these genes might mediate resistance is unclear, as their precise function remains unknown. One possible mechanism may be through mediating the effects of IFN-γ, which orchestrates a plethora of cellular pathways and regulates the expression of hundreds of genes. In mice, IFN-γ driven pathogen resistance is dependent on certain families of GTPases [70,71]. Resistance to *Toxoplasma gondii* requires IFN-γ and it was recently shown that GIMAP proteins mediate resistance to *T. gondii* infection in the resistant Lewis rat strain, with overexpression of GIMAPs in rat macrophages showing that GIMAP 4 had the highest inhibitory effect [72].

### Differences in parasite recognition in the intestine of resistant and susceptible fish

The lack of a transcriptomic response, including any upregulation of immune genes, in the intestine of susceptible fish at 7 dpe was surprising given the high parasite load present in this tissue at that time (Fig 3), and that initial invasion would have occurred 2-3 days prior [9]. This would indicate that susceptible fish are unable to recognize the parasite invading the intestine or the subsequent proliferation. In contrast, resistant fish were able to either prevent parasite establishment in the intestine or minimize parasite proliferation once there. Consistent with this, we observed upregulation of several immune genes in resistant fish. Immunoglobulin kappa constant, which encodes the constant region of immunoglobulin light chains, was mildly upregulated. Fucolectin 6, an F-type lectin that binds fucose was highly upregulated at this timepoint. Lectins are carbohydrate-binding proteins that play a key role in the innate immune response by recognizing exposed glycans on the surface on pathogens [73]. We also observed upregulation of the same homolog of NLRC 5 that was upregulated in the gills of resistant fish at 1 dpe. NOD-, LRR- and CARD-containing (NLRC) proteins are a group of pattern recognition receptors that play a role in both innate and adaptive immune responses by inducing transcription of pro-inflammatory and MHC class I genes, and triggering formation of the “inflammasome”, a multi-protein complex that results in programmed cell death [74,75]. NLRCs are known to play a role in the mucosal immune system of the mammalian gut and are highly expressed by macrophages and epithelial cells in the intestine [76]. Numerous studies of teleost fish have demonstrated the presence of NLRCs that are induced upon immune stimulation or exposure to a pathogen [77–85]. With the generation of several high quality teleost genomes, it is evident that a shared expansion of NLRC genes has occurred in teleosts, suggesting a more prominent role in the immune system [86]. Considering that myxozoans predate the evolution of fish and have been co-evolving with their acquired vertebrate hosts for hundreds of millions of years [87], it seems plausible that fish would have evolved innate immune receptors capable of recognizing conserved motifs on these ubiquitous pathogens.

### Susceptible fish exhibit a vigorous yet ineffective T_H1_ response

Evidence of a strong T_H1_ response was observed in susceptible fish at both 14 and 21 dpe, with upregulation of IFN-γ, its cognate receptor, and T-bet, the master transcriptional regulator of T_H1_ differentiation. GO enrichment analysis also revealed that genes involved in the interferon-gamma signaling pathway were over-represented among the upregulated genes. Upregulation of IFN-γ has been observed in previous studies of Chinook and rainbow trout exposed to *C. shasta* [23,29,31] and appears to play a pivotal and conserved role in the fish response to myxozoan infections. Studies of resistant and susceptible rainbow trout exposed to the myxozoan *Myxobolus cerebralis,* the causative agent of whirling disease, have shown a strong induction of IFN-γ and interferon regulatory factor 1 in both strains, with IFN-γ being upregulated earlier in the infection in resistant fish [88,89]. Olive flounder *(Paralichthys olivaceus)* infected with the myxozoan *Kudoa septempunctata* had elevated levels of IFN-γ in their trunk muscles [90]. IFN-γ was also found to be upregulated in turbot during the early stages of enteromyxosis caused by *E*. *scophthalmi* [91]. Most interestingly, when gilthead sea bream *(Sparus aurata L.)* were exposed to *E. leei,* only the non-parasitized fish had elevated levels of IFN-γ, suggesting it helps mediate resistant to the pathogen [64].

If the IFN-γ pathway is a primary way of defending against myxozoan infections, it raises the question as to why it’s activation in susceptible fish offered no apparent protection against *C. shasta* pathogenesis. Bjork et al. [23] suggest that upregulation of the potent antiinflammatory cytokine IL-10 in susceptible fish may attenuate their inflammatory response and subsequent ability to control parasite proliferation. In concordance with that, we observed marked upregulation of several IL-10 homologs at both timepoints. The ability of IL-10 to attenuate IFN-γ driven parasite clearance by inhibiting the activity of macrophages, T_H1_ cells, and natural killer cells is well-documented [58,92,93]. These immunosuppressive effects are exploited by certain pathogens, including koi herpesvirus, which encodes and expresses a functional IL-10 homolog [94]. Dysregulation of IL-10 production, in terms of timing or overexpression, may explain why susceptible fish fail to inhibit parasite proliferation despite upregulation of IFN-γ.

### The breakdown of the intestinal barrier in susceptible fish

The mucosal surface of the intestine must function as a site of nutrient absorption while acting as a barrier against the systemic spread of microorganisms, both commensal and pathogenic. The main physical component of the intestinal barrier is formed by a continuous monolayer of cells tightly attached to each other by tight junctions, adherens junctions, and desmosomes. Breakdown of this barrier can result in the systemic spread of harmful bacteria and molecules. *C. shasta* reaches the intestine via blood vessels and then migrates through the tissue layers to release spores into the intestinal lumen. As recently shown by Alama-Bermejo et al. [27], *C. shasta* genotype II is highly mobile and has strong adhesive affinities for the glycoprotein components of the extracellular matrix (ECM), resulting in massive interaction and disruption of the host intestinal ECM. We found that genes related to the ECM and cell adhesion showed an intense amount of transcriptional activity in susceptible fish at both 14- and 21 dpe. This aligns with the breakdown of the intestinal structure we observed in histological sections of these fish (Fig 2A-C). Disrupted cell adhesion and cell-to-cell contact also interferes with intercellular communication through gap junctions, which is critical for maintaining tissue structure and homeostasis. Additionally, it can also lead to anoikis, a form of programmed cell death that occurs upon detachment from the ECM. The inability of susceptible fish to overcome *C. shasta* induced breakdown of the ECM would explain why we don’t observe an organized tissue response to the infection (granulomas, fibrosis), as observed in resistant fish.

It is likely that this disruption of the host intestinal barrier and ECM in susceptible fish also lead to the dissemination of bacteria into the intestinal tissue, as evidenced by the upregulation of numerous toll-like receptors that recognize bacterial motifs, as well as cathelicidins, lysozyme, and complement proteins. Pathway level analysis showed the overall immune response transitioned from being primarily IFN-γ driven at 14 dpe (Fig 7A), to a more mixed immune response at 21 dpe (Fig 8A). This likely influx of bacteria coincided with the downregulation of T_H17_ markers IL-17A, IL-17F, and ROR-gamma. T_H17_ cells play a critical role in the response to bacterial pathogens at the gut mucosal surface, and the expression of IL-17A and IL-17F generally increases after exposure to an intestinal pathogen [95–97]. It should also be noted that IL-17F was also downregulated in the gills of susceptible fish at 1 dpe. Whether this represents a maladaptive host response, or a pathogenic strategy remains to be determined. However, it has been shown that certain pathogens actively interfere with the host IL-17 pathway. The mucosal pathogen *Candida albicans* inhibits IL-17 production in human hosts, which is the primary pathway for elimination of the fungus [98], and the intracellular bacteria *Coxiella burnetii* blocks IL-17 signaling in human macrophages [99].

In addition to the likely dissemination of bacteria caused by the breakdown of the intestinal barrier, the hosts ability to acquire nutrients and produce energy became severely compromised. The downregulated genes at 14- and 21 dpe primarily clustered around metabolic and energy producing pathways (Fig 7B, 8B). This occurs while the host is trying to mount a massive immune response, an energetically costly endeavor. This highlights the uphill battle that susceptible fish face: their delayed response to *C. shasta* means they must overcome an evolutionarily well-adapted pathogen that has replicated extensively, while doing so under metabolic stress and with a compromised intestinal structure.

## Conclusions

The primary goal of this study was to determine if susceptible fish recognized *C. shasta* during the initial stages of infection. It is clear from the results at 7 dpe that they fail to recognize the parasite invading the intestine. We specifically used RNA-seq with a high number of replicates to give us the widest possible chance of seeing any genes that respond to the infection, but none were detected. Whether susceptible fish recognize *C. shasta* in the gills remains unclear. We detected a transcriptomic response to the infection; however, this may be actively induced by the parasite and not by host recognition. The observation that both the sympatric (resistant) and allopatric (susceptible) hosts exhibited a similar gill response, and that susceptible fish had no response in the intestine at 7 dpe, supports the idea that the transcriptomic response is driven by the parasite and not by specific host recognition.

The second goal of this study was to identify putative *C. shasta* resistance genes, particularly innate immune receptors that could initiate the immune response. We observed upregulation of a NOD-like receptor whose elevated expression coincided with initial invasion of the gills and intestine. We also observed strong induction of two homologs of GTPase IMAP family member 4 in the gills of resistant fish and later on in the intestine of susceptible fish. Our laboratory is currently in the process of creating a QTL cross of *C. shasta*-resistant and susceptible *O. mykiss* to identify the genomic loci responsible for resistance. Locating these putative resistance genes within the identified loci would offer robust support for their involvement in *C. shasta* resistance and provide a potential marker for rapid identification of resistant fish stocks.

While not an initial goal of this study, we characterized the intestinal response of susceptible fish during the middle and late stages of *C. shasta* infection. As expected from previous studies of *C. shasta* and other myxozoan infections, the immune response was characteristic of an IFN-γ driven T_H1_ response. This response failed to offer any protection though, possibly due to excessive or mistimed expression of IL-10, or the suppression of the T_H17_ response. Comparing the intestinal response of susceptible fish to that of resistant fish with a similar *C. shasta* burden would help answer this, and identify what a successful immune response to the parasite looks like once it has invaded the intestine and begun to replicate.

*C. shasta* is an important pathogen of salmonid fish in the Pacific Northwest and has had an outsized impact on the Klamath River Basin fisheries. As for most myxozoans, what the parasite does within the host and how the host responds has largely remained a black box. The work presented here helps shed light on this process. More broadly, it improves our understanding of myxozoan-host interactions and in conjunction with other studies, may allow general patterns to emerge regarding the fish host’s response. One such pattern may be the conserved adaption of IFN-γ to combat myxozoan infections. This immediately raises the question of how a pathway that is classically associated with the immune response to intracellular pathogens mediates resistance to extracellular myxozoan parasites. Finally, we have identified putative resistance genes that can provide a starting point for future functional studies.

## Supporting information

S1 Table. Primer sequences for qPCR assay

S2 Table. Complete list of differential gene expression results and corresponding GO enrichment.

## Acknowledgments

The authors would like to thank the staff at the John L. Fryer Aquatic Animal Health Laboratory for assistance with fish husbandry, and Dr. Stephen Atkinson and Dr. Rich Holt for assistance with fish sampling. We also thank the Oregon Department of Fish and Wildlife, in particular the Round Butte and Alsea hatcheries for suppling fish used in this study. We would also like to acknowledge Dr. Julie Alexander, Ryan Craig, and Milan Sengthep for their labor-intensive maintenance of the annelid cultures that enable laboratory challenges with *C. shasta*.

## Author Contributions

**Conceptualization:** DB, JB.

**Data curation:** DB.

**Formal analysis:** DB.

**Funding acquisition:** JB.

**Investigation:** DB.

**Methodology:** DB, JB.

**Project administration:** JB.

**Resources:** DB, JB.

**Software:** DB.

**Supervision:** JB.

**Validation:** DB.

**Writing - original draft:** DB.

**Writing - review & editing:** JB.

